# The clinically approved antiviral drug sofosbuvir impairs brazilian zika virus replication

**DOI:** 10.1101/061671

**Authors:** Caroline Q. Sacramento, Gabrielle R. de Melo, Natasha Rocha, Lucas Villas Bôas Hoelz, Milene Mesquita, Caroline S. de Freitas, Natalia Fintelman Rodrigues, Andressa Marttorelli, André C. Ferreiral, Giselle Barbosa-Lima, Mônica M. Bastos, Eduardo de Mello Volotão, Diogo A. Tschoeke, Luciana Leomil, Fernando A. Bozza, Patrícia T. Bozza, Nubia Boechat, Fabiano L. Thompson, Ana M. B. de Filippis, Karin Brüning, Thiago Moreno L. Souza

## Abstract

Zika virus (ZIKV) is a member of *Flaviviridae* family, as other agents of clinical significance, such as dengue (DENV) and hepatitis C (HCV) viruses. ZIKV spread from Africa to Pacific and South American territories, emerging as an etiological pathogen of neurological disorders, during fetal development and in adulthood. Therefore, antiviral drugs able to inhibit ZIKV replication are necessary. Broad spectrum antivirals, such as interferon, ribavirin and favipiravir, are harmful for pregnant animal models and women. The clinically approved uridine nucleotide analog anti-HCV drug, sofosbuvir, has not been affiliated to teratogenicity. Sofosbuvir target the most conserved protein over the members of the *Flaviviridae* family, the viral RNA polymerase. We thus studied ZIKV susceptibility to sofosbovir. We initially characterized a Brazilian ZIKV strain for use in experimental assays. Sofosbuvir inhibits the Brazilian ZIKV replication in a dose-dependent manner, both in BHK-21 cells and SH-Sy5y, by targeting ZIKV RNA polymerase activity, with the involvement of conserved amino acid residues over the members of *Flaviviridae* family. The identification of clinically approved antiviral drugs endowed with anti-ZIKV could reduce the time frame in pre-clinical development. Altogether, our data indicates that sofosbuvir chemical structure is endowed with anti-ZIKV activity.

## INTRODUCTION

Zika virus (ZIKV) is a member of the *Flaviviridae* family, which includes several agents of clinical significance, such as dengue (DENV), hepatitis C (HCV), west Nile (WNV) and Japanese encephalitis (JEV) viruses, among others. This emerging pathogen is an enveloped, positive-sense single stranded RNA virus. Although ZIKV is an arthropod-borne virus (arbovirus) transmitted by mosquitos of the genus *Aedes* (1), transmission through sexual contact have been described (2).

In 1947, in the Zika forest of Uganda, ZIKV was originally identified in sentinel monkeys (3). After occasional episodes of infection in humans in the 50’s, outbreaks have been registered in 2007 (Federated States of Micronesia) and 2013 (French Polynesia) (1). Computational analyses suggest that ZIKV may have been introduced in Brazil already in 2013 (4). In 2015, ZIKV explosively spread across the Brazilian territory and to neighbor countries. Although the epidemiological numbers of zika infection in Brazil may be underestimated – due to limited resources for patient assessments in area where people live below the poverty line – it was predicted that more than 4 million persons were infected (5). ZIKV provokes mild and self-limited exanthematic disease with no or low-grade fever for many patients (6). Remarkably, however, ZIKV infection has been associated to congenital malformations, including microcephaly, and Guillain-Barré syndrome (GBS), based on clinical and laboratorial data (7), (8). Consequently, the World Health Organization (WHO) declared ZIKV infection as a public health emergency of international concern. Antiviral treatments against ZIKV are therefore necessary, because they could not only mitigate ZIKV morbidities but also impair chain of transmission and possess prophylactic activity.

Several broad-spectrum antiviral agents are harmful over pregnant animal models and women. Interferons (IFNs) are abortive, ribavirin and favipiravir are teratogenic (9), (10). Currently, at least three studies have been published in the field of small molecules inhibitors of anti-ZIKV replication (11–13). In these studies, the authors used in their experimental infections the African ZIKV (ZIKV^AFR^) as a prototype (11–13). Importantly, it has been shown that ZIKV^AFR^ is more virulent than the ZIKV strain circulating in Brazil (ZIKV^BRA^) (14), meaning that one might neglected a promising clinically approved compound by screening libraries of compounds over ZIKV^AFR^.

Delvecchio et al. evaluated the pharmacological activity of chloroquine against ZIKV replication (11). Whether chloroquine main mechanism of action relates to the blockage of viral life cycle or promotion of cellular defenses needs to be detailed (11). The studies from Zmurko et al. and Eyer et al. show the anti-ZIKV activity novel nucleoside analogs (12, 13). As these last two works have focused on novel molecules, extensive pre-clinical studies must be carried out before the translation of their data into clinical trials.

Among the *Flaviviridae* family, the gene encoding the RNA polymerase shows the highest degree of conservation (15). Therefore, new therapeutic options against HCV, specially targeting the viral RNA polymerase, could have a broader spectrum over other members of the *Flaviviridae* family. In this regard, sofosbuvir was clinically approved in the last years for therapeutic intervention against HCV infection. Sofosbuvir is a phosphoramidate uridine nucleotide prodrug, which has to be triphosphorylated within the cells to aim the viral RNA polymerase (16). The Food and Drug Administration (FDA) categorizes sofosbuvir as a class B substance: “Animal reproduction studies have failed to demonstrate a risk to the fetus and there are no adequate and well-controlled studies in pregnant women”. The Australian’s equivalent agency, Therapeutic Goods Administration (TGA), suggests a safer use of sofosbuvir, by categorizing this substance as B1: “Drugs which have been taken by only a limited number of pregnant women and women of childbearing age, without an increase in the frequency of malformation or other direct or indirect harmful effects on the human fetus having been observed”. Altogether, these information motivated us to investigate whether sofosbuvir chemical structure possesses anti-ZIKV activity.

## MATERIAL AND METHODS

**Reagents.** The antiviral sofosbuvir, β-d-2’-deoxy-2’-α-fluoro-2’-β-C-methyluridine, was donated by the BMK Consortium: Blanver Farmoquímica Ltda; Microbiológica Química e Farmacêutica Ltda; Karin Bruning & Cia. Ltda, (Toboão da Serra, São Paulo, Brazil). Ribavirin was received as donation form the Instituto de Tecnologia de Farmacos (Farmanguinhos, Fiocruz). Sofosbuvir triphosphate (STP), β-d-2’-deoxy-2’-α-fluoro-2’-β-C-methyluridine triphosphate, ribavirin triphosphate (RTP) and AZT-triphosphate (AZT-TP) were purchased (Codontech.org, CA and Sierra Bioresearch, AZ). Interferon-alpha was purchased from R&D bioscience. All small molecule inhibitors were dissolved in 100 % dimethylsulfoxide (DMSO) and, subsequently, diluted in culture or reaction medium minimally 10^4^-fold before each assay. The final DMSO concentrations showed no cytotoxicity. Materials for cell culture were purchased from Thermo Scientific Life Sciences (Grand Island, NY), unless otherwise mentioned.

**Cells.** Human neuroblastoma (SH-Sy5y; ATCC), baby hamster kidney (BHK-21) and African green monkey kidney cells (Vero) cells were cultured in MEM:F-12 (1:1), MEM and DMEM, respectively. *Aedes albopictus* cells (C6/36) were grown in L-15 medium supplemented with 0.3% tryptose phosphate broth, 0.75 g/L sodium bicarbonate, 1.4 mM glutamine, and nonessential amino acids. The culture medium of the cell types was supplemented with 10 % fetal bovine serum (FBS; HyClone, Logan, Utah), 100 U/mL penicillin, and 100 μg/mL streptomycin(17, 18). Mammals cells were kept at 37 °C in 5% CO_2_, whereas mosquito cells were maintained at 26°C. Passages of SH-sy5y cells included both adherent and non-adherent cells.

**Virus.** ZIKV was isolated from a serum sample of confirmed case from Rio de Janeiro, Brazil. This sample was received and diagnosed by the Reference Laboratory for Flavivirus, Fiocruz, Brazilian Ministry of Health, as part of the surveillance system against arboviruses(3). ZIKV was originally isolated in C6/36 cells, tittered by plaque-forming assay and further passaged at the multiplicity of infection (MOI) of 0.01. Virus passages were performed by inoculating C6/36 cells for 1 h at 26 °C. Next, residual viruses were washed out with phosphate-buffered saline (PBS) and cells were cultured for an additional 9 days. After this period, cells were lysed by freezing and thawing, centrifuged at 1,500 × *g* at 4 °C for 20 min to remove cellular debris.

ZIKV was purified in between fractions of 50 % and 20 % sucrose. Sucrose gradients were made in 40 mL ultracentrifuge tubes (Ultra-clear; Beckman, Fullerton, CA) in PBS without Ca^++^ and Mg^++^ (pH 7.4). Tubes were allowed to stand for 2 h at room temperature. Up to 20 mL of virus was added to each tube and centrifuged in a SW 28 rotor (Beckman) at 10,000 rpm for 4 h at 4 °C. Fractions were collected and assayed for total protein and for virus-induced hemagglutination (HA), with turkey red blood cells (Fitzgerald Industries International, North Acton, MA). Fractions displaying HA activity (≥ 16 UHA/50 μL) were pooled and dialyzed against PBS without Ca^++^ and Mg^++^ (pH 7.4) and 10 % sucrose overnight at 4 °C. Virus pools were filtered through a 0.22-μm membranes (Chemicon, Millipore, Bedford, NY). Infectious virus titers were determined by plaque assay in BHK-21 cells and stored at - 70 °C for further studies.

**Cytotoxicity assay.** Monolayers of 10^4^ BHK-21 or 5 × 10^4^ SH-Sy5y cells in 96-multiwell plates were treated with various concentrations of sofosbuvir or ribavirin, as an additional control, for 5 days. Then, 2,3-Bis-(2-Methoxy-4-Nitro-5-Sulfophenyl)-2*H*-Tetrazolium-5-Carboxanilide (XTT) at 5 mg/ml was added in DMEM in the presence of 0.01 % of N-methyl-dibenzopirazina methyl sulfate (PMS). After incubation for 4 h at 37 °C, plates were read in a spectrophotometer at 492 nm and 620 nm(19). The 50% cytotoxic concentration (CC_50_) was calculated by non-linear regression analysis of the dose-response curves.

**Plaque forming assay.** Monolayers of BHK-21 in 6-well plates were exposed to different dilutions of the supernatant from yield-reduction assays for 1 h at 37 °C. Next, cells were washed with PBS and DMEM containing 1 % FBS and 3 % carboxymethylcellulose (Fluka) (overlay medium) was added to cells. After 5 days at 37 °C, the monolayers were fixed with 10 % formaldehyde in PBS and stained with a 0.1 % solution of crystal violet in 70 % methanol, and the virus titers were calculated by scoring the plaque forming units (PFU).

**Yield-reduction assay.** Monolayers of 10^4^ BHK-21, Vero or 5 × 10^4^ SH-Sy5y cells in 96-multiwell plates were infected with ZIKV at indicated MOIs for 1 h at 37 °C. Cells were washed with PBS to remove residual viruses and various concentrations of sofosbuvir, or interferon-alpha as a positive control, in MEM with 1 % FBS were added. After 24 h, cells were lysed, cellular debris was cleared by centrifugation, and virus titers in the supernatant were determined by PFU/mL. Non-linear regression of the dose-response curves was performed to calculate the inhibitory activity of ZIKV-induced plaque formation by 50 % (EC_50_).

**Preparation of ZIKV RNA polymerase.** ZIKV RNA polymerase was obtained from ZIKV-infected BHK-21 cells. Cells were infected with ZIKV at a MOI of 10 for 24 h, lysed with buffer containing 0.25 M potassium phosphate (pH 7.5), 10 mM 2-mercaptoethanol (2-ME), 1 mM EDTA, 0.5% Triton X-100, 0.5 mM phenylmethane sulfonylfluoride (PMSF) and 20% glycerol, sonicated and centrifuged at 10,000 × *g* for 10 min at 4 °C. The resulting supernatant was further centrifuged at 100,000 × *g* for 90 min at 4 °C and passed through two ion-exchage columns, DEAE- and phopho-cellulose (17).

**RNA polymerase inhibition assay.** ZIKV RNA polymerase inhibition assays was adapted from previous publication (20). The reaction mixture for measurements ZIKV RNA-dependent RNA-polymerase (RDRP) activity was composed of 50 mM hepes (pH 7.3), 0.4 mM of each ribonucleotide (ATP, GTP, CTP and labelled UTP), 0.4 mM dithiothreitol, 3 mM MgCl_2_, 500 ng of ZIKV genomic RNA and cell extracts. ZIKAV RNA was obtained with QIAmp viral RNA mini kit (Qiagen, Duesseldorf, Germany), according to manufacturer instructions, except for the use of RNA carrier. The reaction mixtures were incubated for 1 h at 30 °C in the presence or absence of the drugs. Reactions were stopped with addition of EDTA to make a 10 mM final solution.

The labelled UTP mentioned above represents an equimolar ratio between biotinylated-UTP and digoxigenin-UTP (DIG-UTP) (both from Roche Life Sciences, Basel, Switzerland). Detection of incorporated labeled UTP nucleotides was performed by amplified luminescent proximity homogeneous assay (ALPHA; PerkinElmer, Waltham, MA). In brief, streptavidin-donor and anti-DIG-acceptor beads were incubated with the stopped reaction mixture for 2 h at room temperature. Then, plates containing were read in the EnSpire^®^ multimode plate reader instrument (PerkinElmer). Different types blank controls were used, such as reaction mixtures without cellular extracts and control reaction mixture without inhibitor and beads. In addition, the extract from mock-infected cells was also assayed, to evaluate the presence of RNA-dependent RNA-polymerase activity unrelated to ZIKV. Non-linear regression curves were generated to calculate IC_50_ values for the dose-response effect of the compounds.

**Comparative molecular modeling.** The amino acid sequence encoding of ZIKV RNA polymerase (ZVRP) (UniProtKB ID: B3U3M3) was obtained from the EXPASY proteomic server (21) (http://ca.expasy.org/). The template search was performed at the Blast server (http://blast.ncbi.nlm.nih.gov/Blast.cgi) using the Protein Data Bank (22) (PDB; http://www.pdb.org/pdb/home/home.do) as database and the default options. The T-COFFEE algorithm was used to provide a multiple-alignment between the amino acid sequences of the template proteins and ZVRP. Subsequently, the construction of the SFV-ZVRP complex was performed using MODELLER 9.16 software(23) that employs spatial restriction techniques based on the 3D-template structure. The preliminary model was refined in the same software, using three cycles in the default optimization protocol. Thus, the model structural evaluation was carried out using two independent algorithmsin the SAVES server (http://nihserver.mbi.ucla.edu/SAVES_3/): PROCHECK software(24) (stereochemical quality analysis); and VERIFY 3D(25) (compatibility analysis between the 3D model and its own amino acid sequence, by assigning a structural class based on its location and environment, and by comparing the results with those of crystal structures).

**Metagenomics and genome assembly.** A 0.3 mL of supernatant containing the ZIKV (2 × 10^5^ PFU) was filtered through 0.22 μm filters to remove residual culture cells. The virus RNA was extracted using QIAamp Viral RNA Mini Kit (Qiagen^®^) with DNAse RNAse-free (Qiagen^®^) treatment, omitting carrier RNA. Double-stranded cDNA libraries were constructed by Truseq Stranded total RNA LT (Illumina^®^) with Ribo-zero treatment, according to the manufacture’s instruction. The library size distribution was assessed using 2100 Bioanalyzer (Agilent^®^) with High Sensitive DNA kit (Agilent^®^), and the quantification was performed with 7500 Real-time PCR System (Applied Biosystems_®_) with KAPA Library Quantification Kit (Kapa Biosystems). Paired-end sequencing (2 × 300 bp) was done with MiSeq Reagent kit v3 (Illumina^®^). The sequences obtained were preprocessed using the PRINSEQ software to remove reads smaller than 50 bp and sequences with scores of lower quality than a Phred quality score of 20. Paired-End reAd merger (PEAR) software was used to merge and extend the paired-end Illumina reads using the default parameters(26), (27). The extended reads were analyzed with Deconseq program, against the Human Genome Database, with Identity and Coverage cutoff of 70%, to remove human RNA sequences(28). Nonhuman reads were analyzed against all GenBank viral genomes (65 □ 052 sequences) trough BLAST software using 1e-5 e-value cutoff. The sequences rendering a genome were assembled with SPAdes 3.7.1 software(29) followed by a reassemble with CAP3 program(30).

**Sequence comparisons.** Sequences encoding for the C-terminal portion of the RNA polymerase form members of the *Flaviviridae* family were acquired from complete sequence deposited in GenBank. Alignment was made using the ClustalW algorithm in Mega 6.0 software. Sequences were analyzed using neighbor-joining, with pairwise deletion, with bootstrap of 1,000 replicated and *P* distances were registered. Sequences were also analyzed for mean evolutionary rate.

**Statistical analysis.** All assays were performed and codified by one professional. Subsequently, a different professional analyzed the results before identification of the experimental groups. This was done to keep the pharmacological assays as blind. All experiments were carried out at least three intendent times, including technical replicates in each assay. The dose-response curves to calculate EC_50_ and CC_50_ values were generated by Excel for Windows. Dose-response curve to calculate IC50 values were obtained by Prism graphpad software 5.0. The equations to fit the best curve were considered based on the R^2^ values ≥ 0.9. Above mentioned are the statistical analysis specific to each program software used in the bioinformatics analysis.

## RESULTS

**Sofosbuvir fits on the ZVRP predicted structure.** The RNA polymerase structures from WNV (PDB # 2HFZ) (31), JEV (PDB # 4K6M) (32), DENV (PDB # 5DTO) (33) and HCV (PDB # 4WTG) (34) share 72, 70, 68, and 25 % sequence identity with ZIKV orthologue enzyme, respectively. Despite that, the HCV enzyme is complexed with sofosbuvir and the amino acids residues that interacts with this drug are highly
conserved over members of the *Flaviviridae* family, around 80 % (34). The region encoding for the C-terminal portion of *Flaviviridae* RNA polymerase contains around the 800 amino acid residues and identical residues are highlighted in yellow (Supplementary Material 1). The residues critical for RDRP activity are conserved among different viral species and strains, including: ZIKV African strain from the 50’s and those circulating currently, DENV and different genotypes of HCV (Supplementary Material 1) (35).

Based on the HCV RNA-direct RNA-polymerase domain, we constructed a 3D model for ZIKV orthologue enzyme (Figure 1). Sofosbuvir was located among the palm and fingers region of ZIKV RNA polymerase (Figure 1A), an area important to coordinate the incorporation of incoming nucleotides into the new strand of RNA (34). Consequently, amino acid residues relevant to sofosbuvir interaction are some of those critical for natural nucleotide incorporation and thus RDRP activity (Figure 1B) (34). The amino acid residues involved with the interaction with sofosbuvir are identical or conserved among the members of the *Flaviviridae* family (Supplementary Material 2). The conserved residues are considered to be evolving slower than other residues of this enzyme (Supplementary Material 2). These information mean that residues predicted to be required for interaction with sofosbuvir tend to be conserved among members of *Flaviviridae* family.

**Figure 1.**
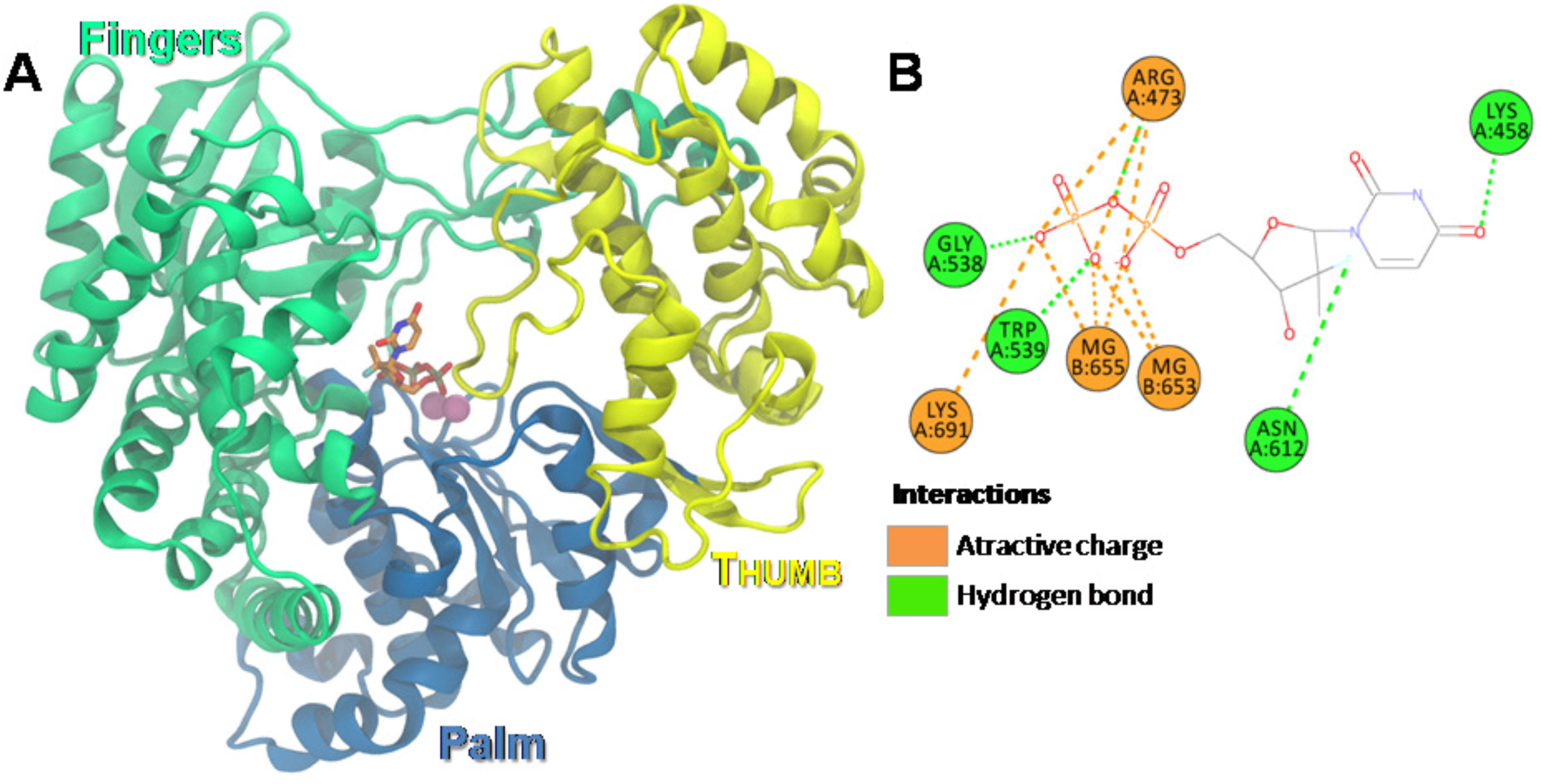
Putative ZIKV RNA polymerase in complex with sofosbuvir. Based on the crystal structure of the HCV RNA polymerase in complex with sofosbuvir diphosphate (PDB accession # 4WTG), the putative structure from ZIKV RNA polymerase was constructed. The amino acid sequence encoding ZIKV RNA polymerase (ZVRP) (UniProtKB ID: B3U3M3) was aligned using T-COFFEE server with orthologue RNA polymerases from members of *Flaviviridae* family, Hepatites C virus (HCV; PDB # 4WTG, West Nile virus (WNIV; PDB # 2HFZ), Japanese Encephalitis virus (JEV; PDB # 4K6M), and Dengue virus (DENV; PDB # 5DTO). The MODELLER 9.16 software was used to build a 3D-model of ZIKV RNA polymerase, with subsequent refinement using three cycles in the default optimization protocol. The model structural evaluation was carried out using two independent algorithms, PROCHECK software and VERIFY 3D. (A) The 3D-model of ZIKV RNA polymerase is presented. (B) The residues presumably required for ZIKV RNA polymerase interaction with sofosbuvir and Mg^++^ ions.

**Sofosbuvir inhibits ZVRP in a dose-dependent fashion.** Next, we evaluated whether sofosbuvir triphosphate (STP), the bioactive compound, could inhibit ZIKV RDRP activity. Fractions containing the ZIKV RDRP activity were purified from infected cells (17). STP inhibited ZIKV RDRP activity with an IC_50_ value of 0.38 ± 0.03 μM (Figure 2). Ribavirin-triphosphate (RTP) and AZT-TP were used as positive and negative controls, respectively (Figure 2). RTP and AZT-TP presented IC_50_ values of 0.21 ± 0.06 and > 10 μM, respectively (Figure 2). The data from Figure 2, confirmed the molecular modeling prediction that sofosbuvir docked onto ZVRP structure and, reveled that sofosbuvir chemical structure inhibits ZIKV RDRP activity.

**Figure 2.**
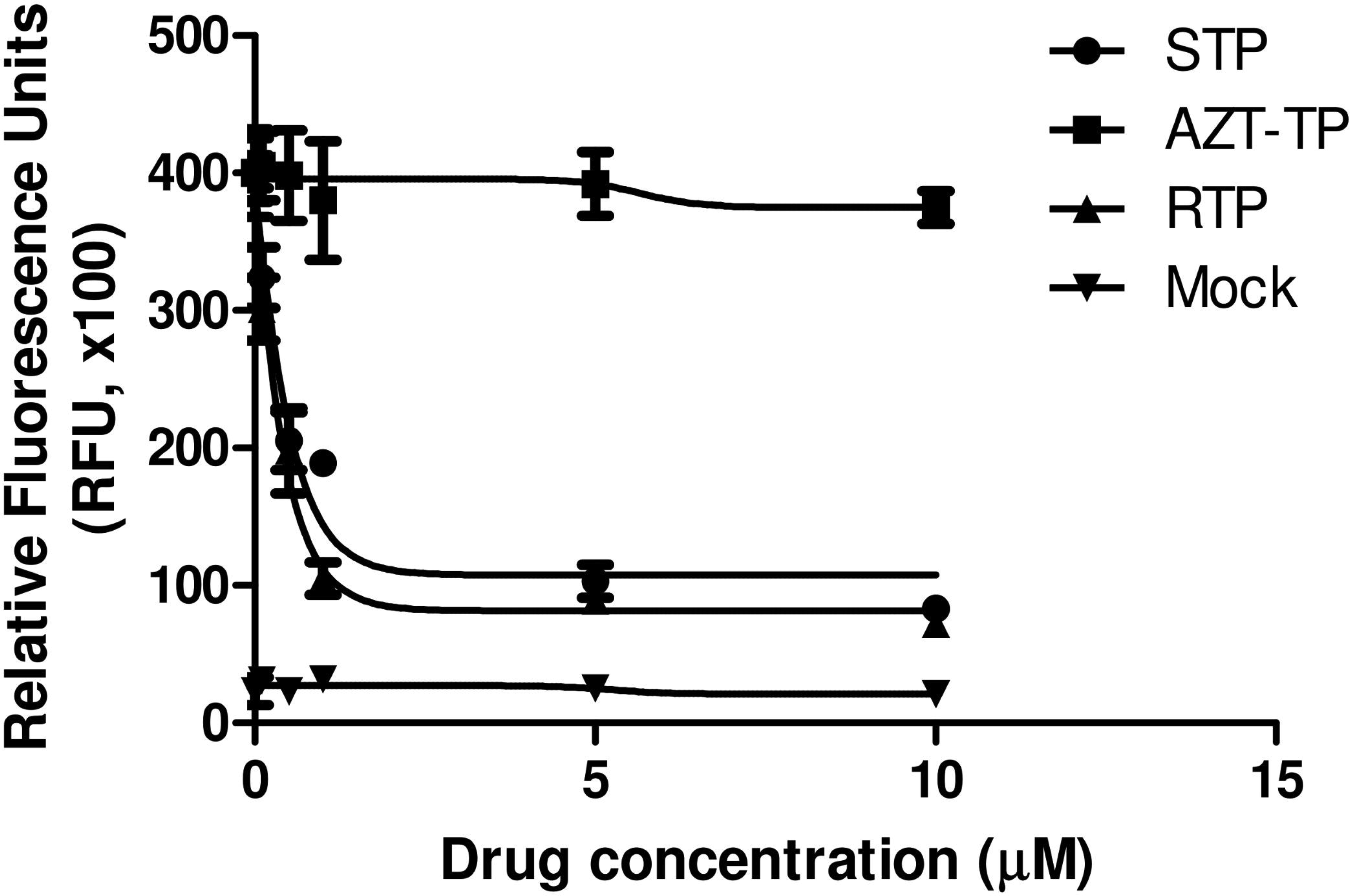
Sofosbuvir inhibits ZIKV RDRP activity. Cell extracts from ZIKV-infected cells were assayed for RDRP activity using virus RNA as the template and labelled UTP as the tracer. biotinylated-UTP and digoxigenin-UTP were detected by ALPHA technology using the EnSpire^®^ multimode plate reader instrument (PerkinElmer). The molecules used assayed were sofosbuvir triphosphate (STP), ribavirin triphosphate (RTP) and AZT triphosphate (AZT-TP). As a control, RNA-dependent RNA-polymerase activity was measured in extracts from mock-infected cells (mock). Data represent means ± SEM of five independent experiments.

**Sofosbuvir inhibits ZIKV replication in cell-, MOI- and dose-dependent manner.** Sofosbuvir phosphoramidate prodrug must be converted to its triphosphate analog within the cellular environment to become active. Therefore, we investigated whether sofosbuvir inhibits ZIKV replication in cellular systems. Before that, we isolated a Brazilian ZIKV strain from a confirmed case of zika fever and characterized this isolate for experimental use. The full-length virus genome was sequenced (GenBank accession # KX197205) and characteristic plaque forming units (PFU) and cytopathic effect (CPE) were detected in BHK-21 cells (Figure S1). Another concern was to stablish whether other plaque-forming viral agents was co-isolated, which could be misleading to interpret the antiviral activity. Metagenomic analysis reveal that ZIKV was the only full-length genome of a plaque-forming virus in BHK-21 detected (Supplementary Material 3).

After the characterization of a Brazilian ZIKV strain for experimental virology assays, we evaluated the ZIKV susceptibility to sofosbuvir. BHK-21, Vero or human neuroblastoma (SH-Sy5y) cells were inoculated at different MOIs and treated with various concentrations of sofosbuvir. Supernatant from these cells were collected and infectious virus progeny tittered. Sofosbuvir induced a MOI-and dose-dependent inhibition of ZIKV replication (Figure 3A and Figure 3B, Table 1 and Figure S2). Potency and efficiency to inhibit ZIKV replication were higher in SH-Sy5y than BHK-21 cells (Figure 3A and Figure 3B, Table 1 and Figure S2). Of note, over 10 μM of sofosbuvir did not inhibit ZIKV replication in Vero cells, indicating a cell-depend inhibition of ZIKV replication. IFN-alpha and ribavirin were used as positive controls to inhibit ZIKV replication (Figure 3A and Figure 3B, Table S1 and Figure S2).

**Figure 3.**
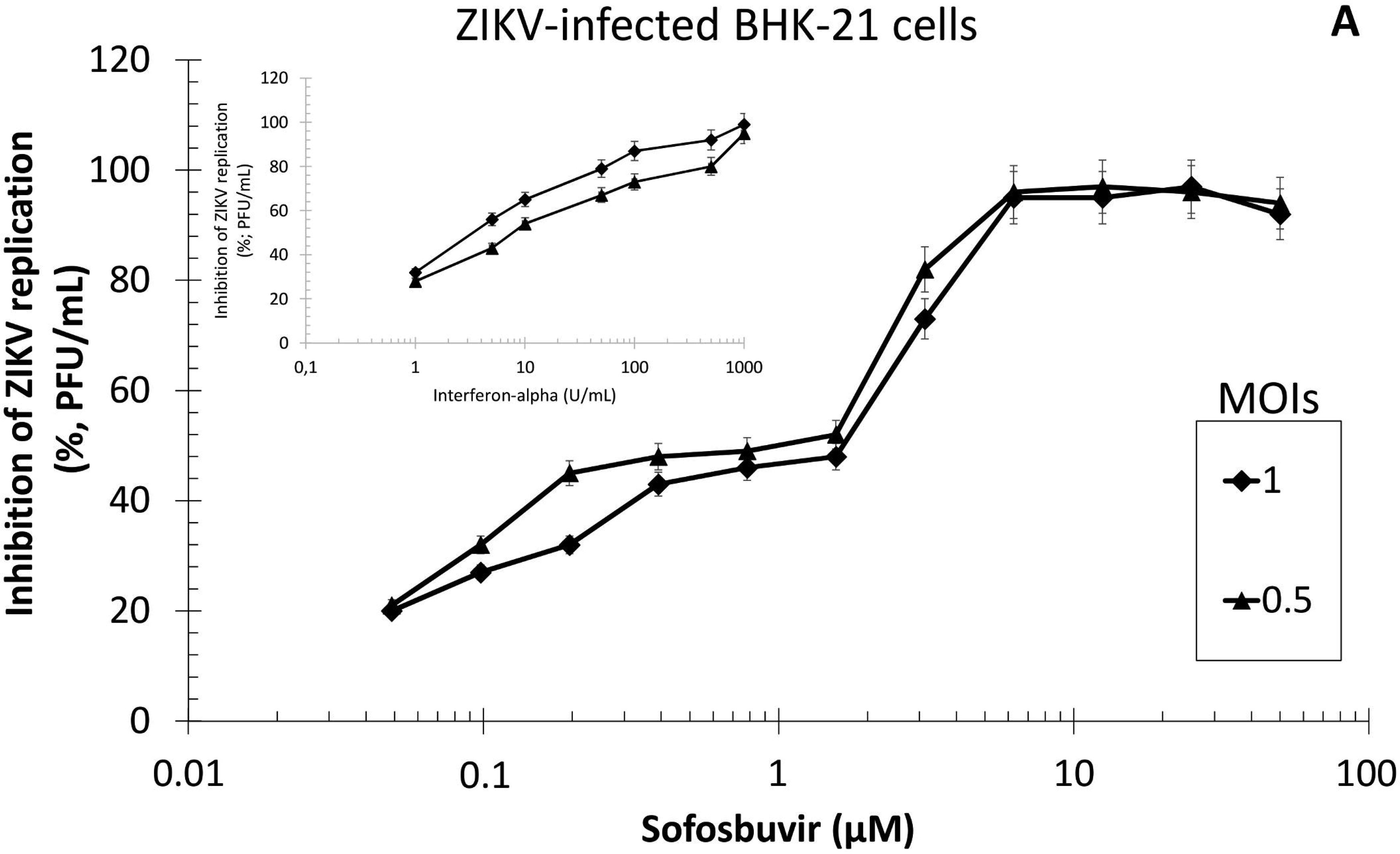
The antiviral activity of sofosbuvir against ZIKV. BHK-21 (A) or SH-Sy5y (B) cells were infected with ZIKV at indicated MOIs, exposed to various concentrations of sofosbuvir or IFN-alpha (inset), and viral replication was measured by plaque-forming assay after 24 h of infection. Data represent means ± SEM of five independent experiments.

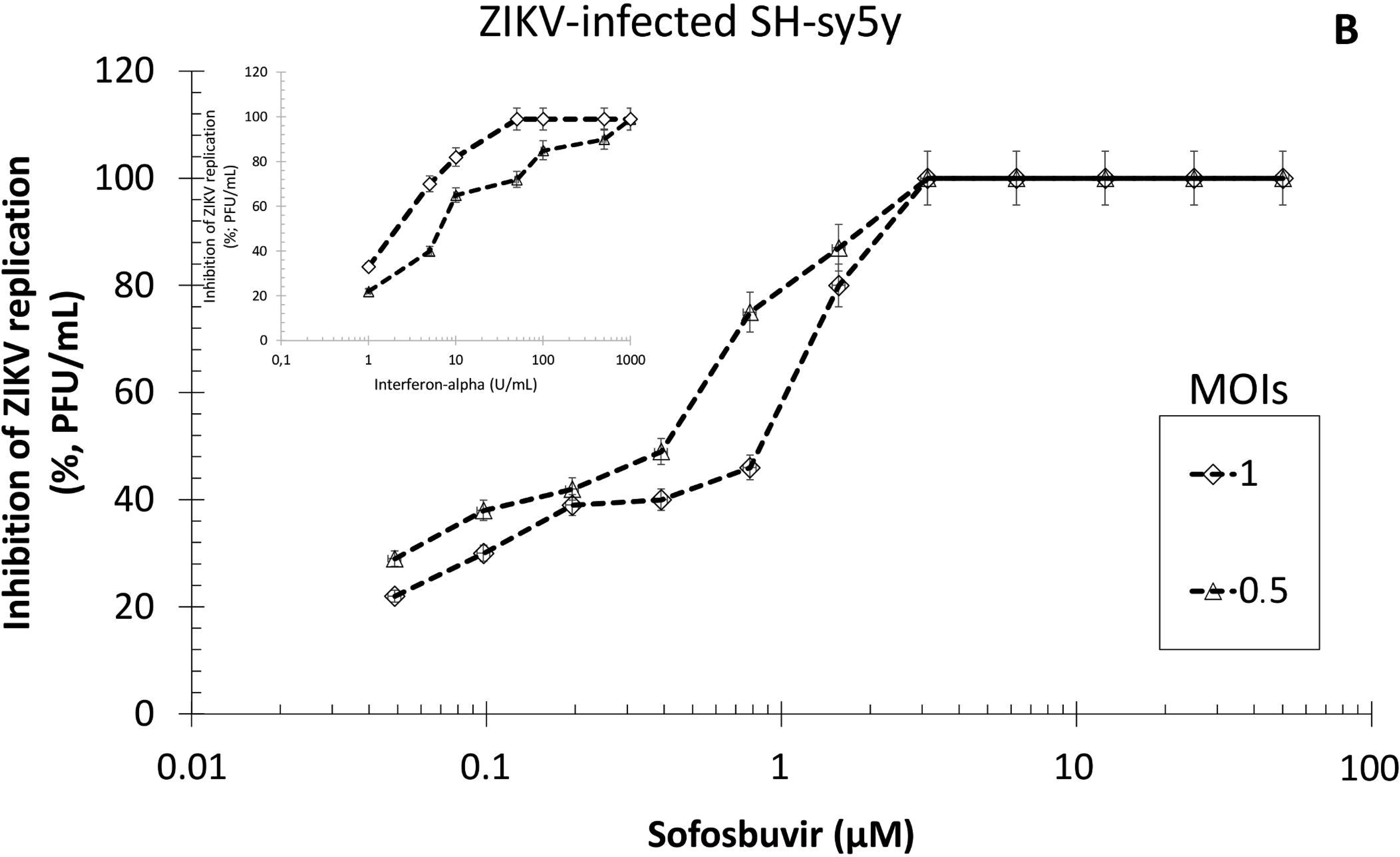

**Table 1.** Antiviral activity and cytotoxicity of sofosbuvir.

| Pharmacological |  |  |  |  |  |  |  |  |  |  |  |  |  |  |  |
| --- | --- | --- | --- | --- | --- | --- | --- | --- | --- | --- | --- | --- | --- | --- | --- |
| Parameter |  | EC<sub>50</sub> (μM) |  |  |  |  | CC<sub>50</sub> (μM) |  |  | SI |  |  |  |  |  |
| Cell type | BHK |  | SH-Sy5Y |  | Vero |  | BHK | SH-Sy5Y | Vero | BHK |  | SH-Sy5Y |  | Vero |  |
| MOI | 1.0 | 0.5 | 1.0 | 0.5 | 1.0 | 0.5 |  |  |  | 1.0 | 0.5 | 1.0 | 0.5 | 1.0 | 0.5 |
| Sofosbuvir | 1.9 ± 0.2 | 1.7 ± 0.1 | 1.1 ± 0.2 | 0.65 ± 0.08 | >10 | >10 | 360 ± 43 | 421 ± 34 | 461 ± 32 | 184 | 212 | 384 | 653 | NA | NA |
| Ribavirin | 5.3 ± 0.8 | 3.1 ± 0.6 | 2.9 ± 0.4 | 1.2 ± 0.2 | 6.8 ± 1.2 | 4.8 ± 0.8 | 177 ± 22 | 300 ± 21 | 371 ± 16 | 33 | 57 | 103 | 250 | 54 | 77 |
| IFN-alpha* | 7.3 ± 0.3 | 2.8 ± 0.3 | 9.8 ± 1.2 | 4.9 ± 0.6 | 10.2 ± 1.6 | 5.9 ± 1.0 | ND | ND | ND | NA | NA | NA | NA | NA | NA |

\*Values are expressed as U/ml, ND – Not determined, NA – Not Applicable

Sofosbuvir cytotoxicity was also cell type-dependent (Table 1), being less cytotoxic for SH-Sy5y than for BHK-21 cell line. Our results indicate that the selectivity index (SI; which represents the ratio between CC_50_ and EC_50_ values) for sofosbuvir varied from 185 to 653 (Table 1) – being safer at MOI equals to 0.5 in the neuroblastoma cell line. For comparisons, SI values for sofosbuvir were up to 5 times higher than for ribavirin (Table 1). Our data point out that that sofosbuvir chemical structure is endowed with anti-ZIKV activity and efforts to promote its broader activation in different cellular types could lead to the development to new anti-ZIKV based therapies.

## Discussion

ZIKV is a member of *Flaviviridae* family, such as other clinically relevant viruses, such as DENV, WNV, JEV and HCV. Among this family, ZIKV was considered to be a virus causing only mild and self-limited infections (6). However, based on clinical evidence and laboratorial data, ZIKV infection was associated with neurological-related morbidities, with impacts on the development of human nervous system and triggering of GBS (7), (8, 14), (36–39). Antiviral treatment options are thus required to block viral replication. Previous studies have demonstrated that the anti-malarial drug chloroquine and novel nucleoside analogs inhibit ZIKV replication (11–13). Here, we show that the clinically approved uridine nucleotide analog anti-HCV drug, sofosbuvir, is endowed with anti-ZIKV activity.

Delvecchio et al. evaluated the pharmacological activity of chloroquine against ZIKV replication (11). The authors used a clinically approved drug feasible to be used in pregnant women. The pharmacological activity of chloroquine against ZIKV replication in Vero cells, which produce high virus titers, and relevant cellular models for studying ZIKV neurotropic replication were evaluated. The African ZIKV isolate used, which is differently than the one circulating currently. Besides, chloroquine’s potency over ZIKV replication is around 10 μM, whereas against different species of *Plasmodim* it acts at the sub-micromolar range. This means that the 500 mg chloroquine tablets approved for clinical use to treat malaria might not be enough to treat ZIKV infection. The mechanism by which chloroquine inhibits ZIKV replication has not been characterized. Whether it inhibits the viral life cycle directly or enhances cell survival must be investigated.

The studies from Zmurko et al. and Eyer et al. show the anti-ZIKV activity novel nucleoside analogs (12), (13). Zmurko et al. characterized their studied molecule *in vitro*, indicating its ability to target the ZIKV RNA polymerase, and *in vivo* (12). Eyer et al. evaluated a broad range of nucleoside/nucleotide analogs, including clinically approved drugs, pointing out to another novel molecule (13). In these studies, the authors also used the African ZIKV strain and the pharmacological activity has not been characterized in neuronal cell lines. As these works have focused on novel molecules, the translation of their data into clinical trials will require further extensive studies.

Sofosbuvir chemical structure is endowed with anti-ZIKV activity, by different approaches we observed that. Predicted ZVRP structure suggest that sofosbuvir required critical amino acid residues for ribonucleotide incorporation, such as Arg473, Gly538, Trp539, Lys691. These results could anticipate that genetic barrier for antiviral resistance would be high. That is, changes in these residues would jeopardize enzyme activity. Such a high genetic barrier is found to the emergence of sofosbuvir-resitant strains of HCV (34). The fluoride radical in sofosbuvir rybosil moiety is coordinated by the Asn612, an interaction involved with the drug selectivity to RDRP, which may avoid unspecific effects towards the cellular DNA-dependent RNA-polymerase. The Lys458 seems to be the docking residue for the uridine analog. Further enzymatic studies with site-directed mutagenesis to this residue could confirm its participation for sofsobuvir docking.

Sofosbuvir produced a dose-dependent inhibition of ZIKV replication with different magnitudes in terms of potency and efficiency in BHK-21 and SH-Sy5y cells. On the other hand, we observed no inhibition of viral replication with over 10 μM in Vero cells. Similarly, in the recent study from Eyer et al. (13), African ZIKV susceptibility sofosbuvir was screened in Vero cells, and this compound did not emerged as a potential hit. Although Eyer et al. (13) and us used different viral strains, we reached similar results in Vero cells. Interestingly, sofosbuvir is a substrate for glycoprotein-P (40). Differently than in BHK-21 and Sh-Sy5y, proteomic data reveal that Vero cells express this multi-drug resistance ABC-transporter, which may cause sofosbuvir efflux out of the cell (41-43).

Of note, although we succeed to determine the sofosbuvir antiviral activity against ZIKV in different cell types, this drug metabolism *in vivo* occurs mainly in the liver and adjacent organs. Different experimental *in vivo* infection assays have been studied (ref). Prior to the detection of the ZIKV in the nervous system, virus is found in peripheral organs, such as in spleen and liver (12), (44-46). We believe this could be an insight from the natural history of the ZIKV infection in patients. That is, it is likely that before reaching sites of immune privilege, such as the nervous system or the placental barrier, ZIKV virions may be amplified in peripheral organs and provoke an inflammation response to disrupt the disrupt specific barriers. Although the role of the liver during ZIKV infection is currently overlooked, ZIKV could replicate in the liver before its systemic spread. Indeed, it has been reported for some ZIKV-infected patients that liver transaminase levels are enhanced during the onset of illness (47). Moreover, DENV viral loads are increased in the liver and represent one of the hallmarks of this other flavivirus pathogenesis. Naturally, liver is the main site of HCV replication. Therefore, *in vivo* experimental assays are necessary to better determine sofosbuvir capacity to acting as a therapeutic or prophylactic agent during ZIKV experimental infection.

ZIKV-associated microcephaly and GBS highlights that antiviral interventions are urgent. Our data reveal that a clinically approved drug is endowed with antiviral activity against ZIKV. Thus, the potential second use of sofosbuvir, an anti-HCV drug, against ZIKV seems to be plausible.

## AUTHORS CONTRIBUTIONS

C.Q.S., G.R.deM., N.R., L.V.B.H., M.M., C.S.deF., N.F.-R., A.M., A C., G.B.-L., E.de M.V., D.A.T. and L.L. - contributed with experiment execution and analysis.

M.M.B., F.A.B., P.B., N.B., F.L.T., A.M.B.deF., K.B. and T.M.L.S. - Data analysis manuscript preparation and revision K.B. and T.M.L.S. - Conceptualized the study All authors revised and approved the manuscript.

## ACKNOWLEDGEMENTS

Thanks are due to Drs. Carlos M. Morel, Marcio L. Rodrigues, Renata Curi and Fabrícia Pimenta for strong advisement and support regarding technological development regarding this project. This work was supported by Conselho Nacional de Desenvolvimento e Pesquisa (CNPq), Fundação de Amparo à Pesquisa do Estado do Rio de Janeiro (FAPERJ).

**The authors declare no competing financial interests.**

## Supplemental Information

**Figure S1.**
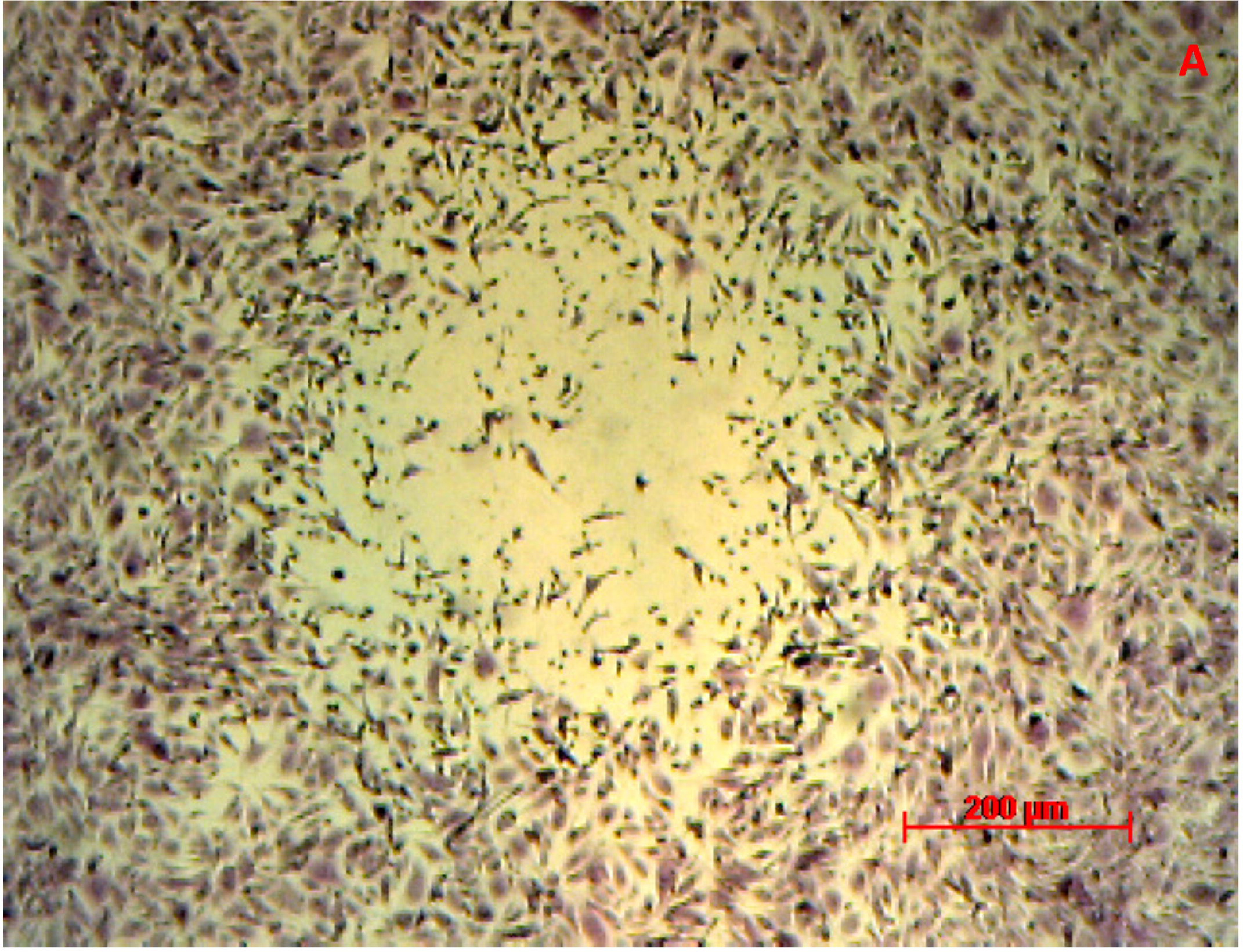
Plaque assay for ZIKV. Monolayers of BHK-21 cells were infected for 1 h at 37 °C. After that, viruses were washed out with PBS and wells were covered with overlay medium with 1 % FBS. At 5 days post-infection, plaques were fixed and colored with crystal violet. (A) A representative plaque forming unit (PFU) is presented at 40 x magnification. (B) The PFU and adjacent cellular monolayer is presented at 100 x magnification, some of these cells present ZIKV-induced cytopathic effect (CPE), three examples are highlighted by the red arrows. (C) A representative closer view of ZIKV-induced CPE (red arrow).

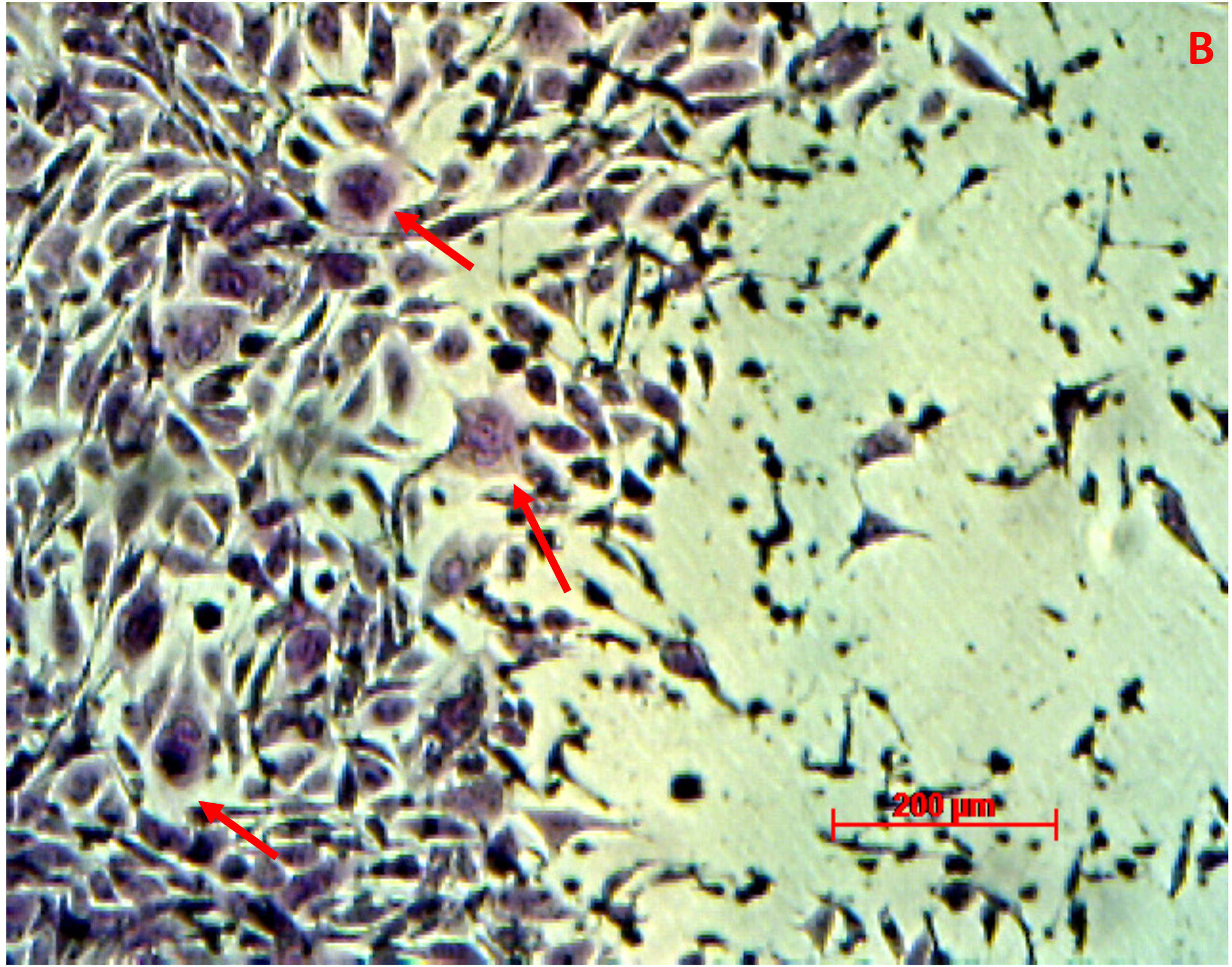

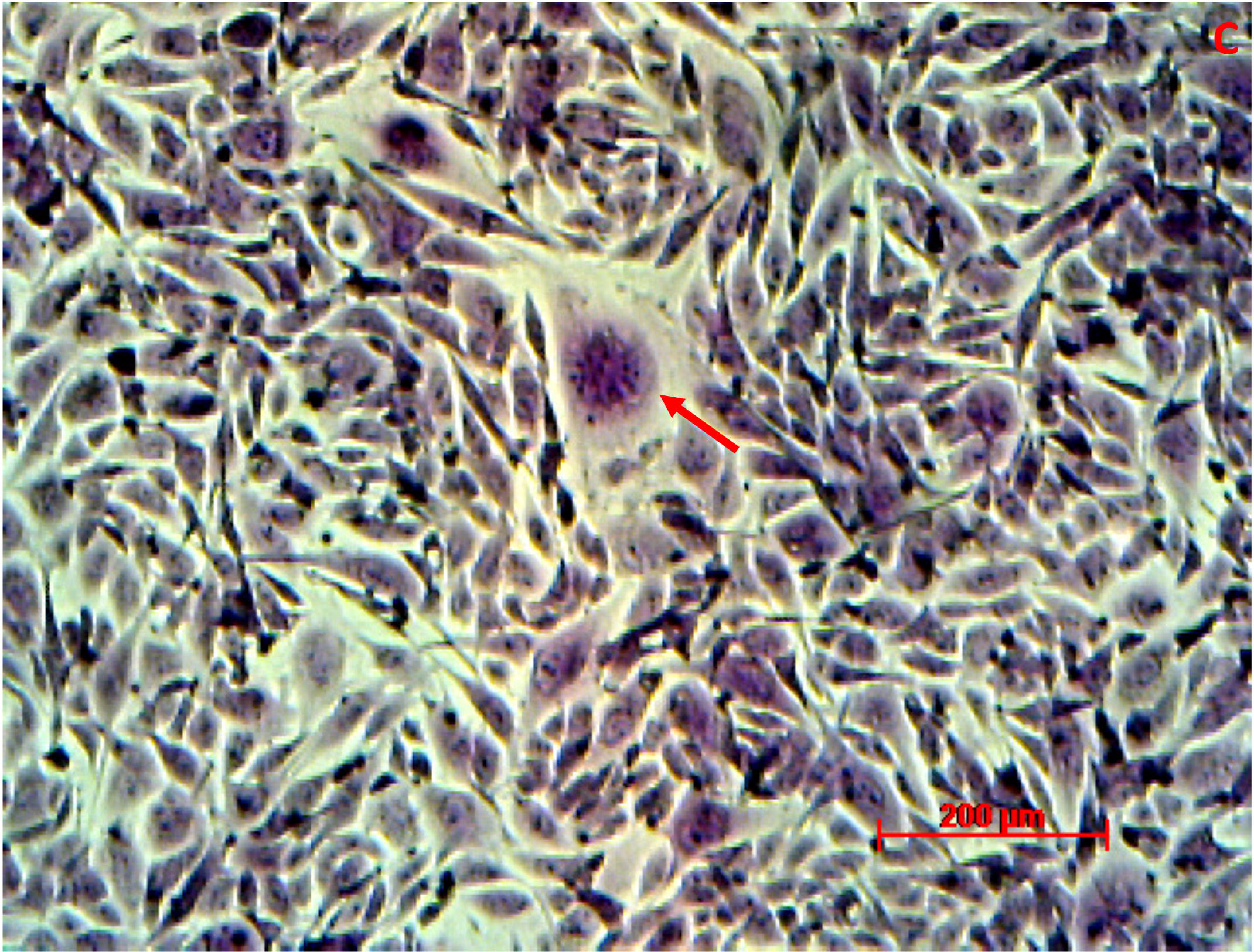

**Figure S2.**
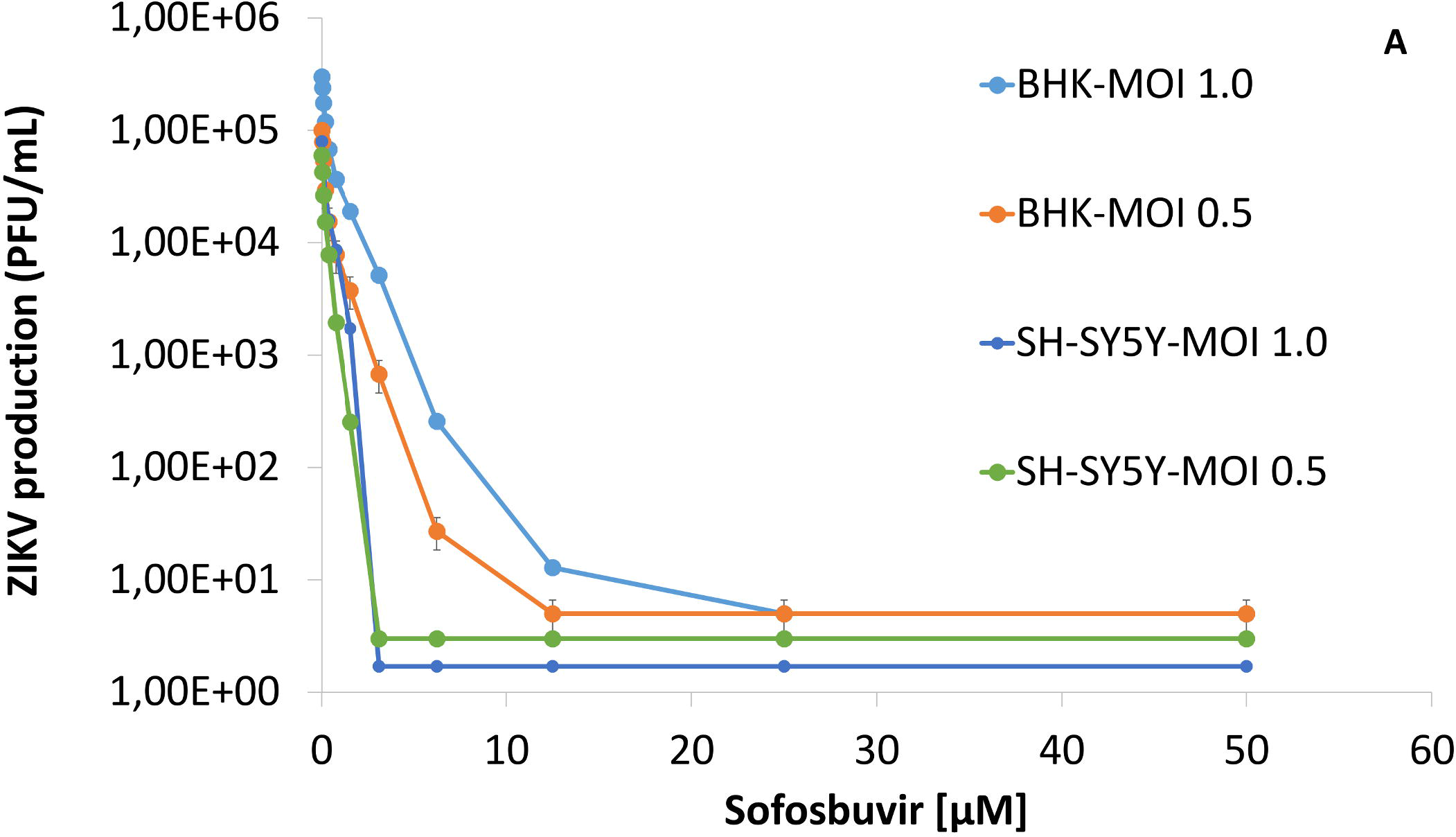
The antiviral activity of Sofosbuvir against ZIKV. BHK-21 or SH-sy5y were infected with ZIKV at indicated MOIs, exposed to various concentrations of sofosbuvir (A) or IFN-alpha (B), and viral replication was measured by plaque-forming assay after 24 h of infection. Data represent means ± SEM of three independent experiments

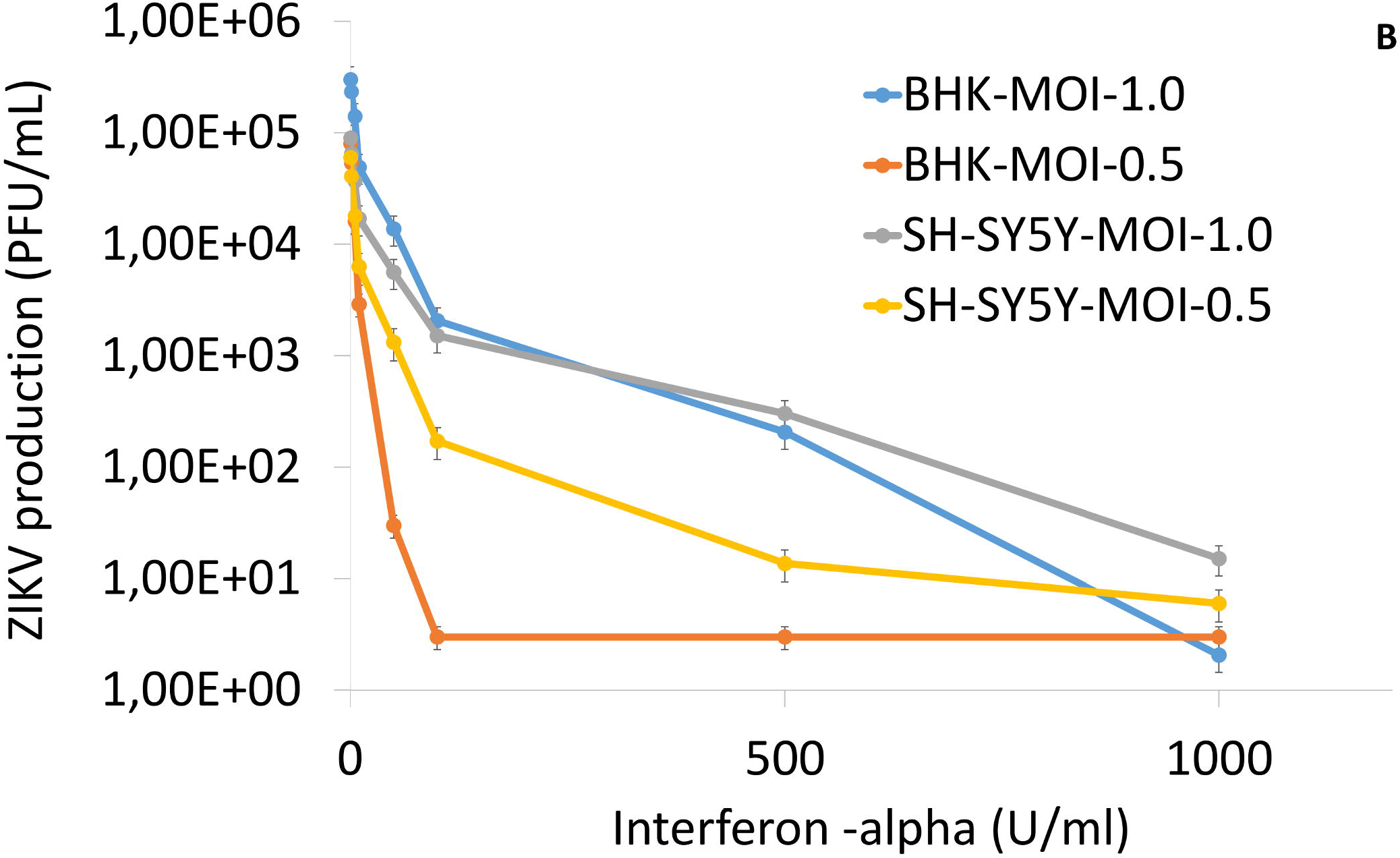

